# Inertial sensor-based centripetal acceleration as a correlate for lateral margin of stability during walking and turning

**DOI:** 10.1101/768192

**Authors:** Peter C. Fino, Fay B. Horak, Carolin Curtze

## Abstract

There is growing interest in using inertial sensors to continuously monitor gait during free-living mobility. Inertial sensors can provide many gait measures, but they struggle to capture the spatial stability of the center-of-mass due to limitations estimating sensor-to-sensor distance. While the margin of stability (MoS) is an established outcome describing the instantaneous mechanical stability of gait relating to fall-risk, methods to estimate the MoS from inertial sensors have been lacking. Here, we developed and tested a framework, based on centripetal acceleration, to determine a correlate for the lateral MoS using inertial sensors during walking with or without turning. Using three synchronized sensors located bilaterally on the feet and lumbar spine, the average centripetal acceleration over the subsequent step can be used as a correlate for lateral MoS. Relying only on a single sensor on the lumbar spine yielded similar results if the stance foot can be determined from other means. Additionally, the centripetal acceleration correlate of lateral MoS demonstrates clear differences between walking and turning, inside and outside turning limbs, and speed. While limitations and assumptions need to be considered when implemented in practice, this method presents a novel correlate for the lateral MoS during walking and turning using inertial sensors, although further validation is required for other activities and populations.

## I. INTRODUCTION

Recent advances in wearable sensors have enabled biomechanical analyses of gait outside of the laboratory. Continuous monitoring of gait during free-living daily activity provides a new window into community ambulation and presents a promising avenue for future gait research investigating older adults at risk of falls [1], neuropathological progression [2], and ecologically valid gait assessments [3–7].

Many spatiotemporal gait parameters, including measures of pace, rhythm, variability, and asymmetry, can be estimated using inertial sensors [4, 8], but spatial stability has been difficult to assess using inertial sensors alone. Inertial sensors can also assess dynamic, temporal stability, derived from short-term maximum finite-time Lyapunov exponents or other dynamical systems constructs, that describe the temporal stability of a system within a given state space [9]. While temporal measures are theoretically valid and predictive metrics of the probability of falling [1, 9], they do not describe the instantaneous biomechanical stability during locomotion.

To describe the mechanical stability of gait, Hof and colleagues proposed extending the inverted pendulum model of human balance using the velocity of the center of mass (CoM) to extrapolate the velocity-adjusted position of the CoM (XcoM)[10, 11]. The relationship between the XcoM and the base of support (BoS) reveals the instantaneous mechanical stability of the system; if the XcoM falls outside the BoS, balance cannot be recovered with ankle joint torque alone – a stepping response, rotation at superior joints, or external force is required[11]. Since the spatial distance between the XcoM and the BoS was defined as the margin of stability (MoS) [10, 11], the MoS has been widely used to assess gait stability [9, 12–17], and gait controllers have been proposed with objectives of maintaining constant MoS through foot placement [18, 19].

Traditionally, MoS has been assessed using optical motion capture, gait carpets, and / or force platforms that give accurate spatial information [16, 20–22]. Inertial sensors, comparatively, provide accurate acceleration, angular velocity, and orientation estimates, but struggle to provide accurate positional distances from one sensor to another. To rectify this issue, static calibration poses and subject-specific anthropometric dimensions have been used to establish initial positions of each sensor [23, 24]. However, requiring the subject to hold a neutral pose for calibration before every data capture may not be a viable solution for continuous monitoring in free-living conditions. Recently, a custom combination of inertial sensors and pressure-sensitive insoles have been used to estimate the position of the CoM [25] and the MoS during walking [26]. While this system provides promise for assessments of MoS during walking outside the laboratory, it relies on custom shoes and may not be feasible for large-scale or long-term monitoring.

Others have described a method of estimating the MoS using a series of inertial sensors on the feet, shank, leg, and hip to reconstruct the kinematic chain [27]. This method achieves low error (<2 cm), but relies on subject-specific anthropometry and either external reference frames or static and dynamic calibration poses for sensor-to-segment calibration. As the authors combined inertial sensor data with motion camera data to establish a global reference frame, it is unclear if the same accuracy can be obtained using inertial sensor data alone [27].

Recently, the lateral MoS was estimated in individuals with dementia within a community-dwelling setting using camera-based systems [15]. While other spatiotemporal gait outcomes were assessed, the estimated lateral MoS was the only gait measure associated with prospective falls [15]. These results support quantifying the lateral MoS in community settings to assess fall-risk. Yet, the novel, low-cost camera system relies on line of sight and cannot assess MoS in every environment. In contrast, inertial sensors are wearable, capable of continuously quantifying ambulation in diverse environments, and becoming a predominant method for community-based assessments. Yet, there are few methods that can quantify MoS without cameras, and there are no established method to estimate MoS using only a single inertial sensor.

Because inertial sensors struggle to provide inter-sensor positional data, we sought to create a framework by which lateral MoS could be easily inferred from acceleration data. Thus, our aim was to develop and validate a correlate for lateral MoS during walking and turning using inertial sensors, with the eventual goal of using a single inertial sensor. As a number of studies and publicly available datasets have utilized a single inertial sensor on the lumbar spine [7, 28], and this location is in close proximity to the whole-body center-of-mass, we focused on using this lumbar-spine location. We compared our correlate of MoS using the inertial sensor to the true MoS based on optical motion capture. To extend the comparison to include a variety of daily ambulatory tasks, we included steps during straight gait and a variety of different turning angles.

## II. METHODS

### A. Model Framework

Based on Hof et al. [10], dynamic balance can be achieved by placing the foot, and the CoP by extension, some offset outside of the XcoM to generate a corrective torque. The instantaneous mediolateral **MoS**_**z**_ is the difference between CoP, ***u***_***z***_, and the **XcoM** based on

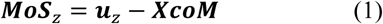

where the **XcoM** is defined by the lateral position of the CoM, ***z***, the lateral velocity of the CoM, ***v***_***z***_, and the eigenfrequency of the inverted pendulum 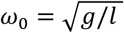 by

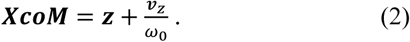

Based on this model and assuming the minimum occurs at or near initial contact [10], several simple controllers can be derived for control of forward and lateral foot placement based on maintaining a minimum MoS at each step, ***MoS***_***zn***_ [10, 19]. For lateral controllers with constant step time, a change in lateral CoM velocity, Δ***v***_***z***_, occurring over the previous step *n-1* can be corrected through a change in foot position of the current step *n* equal to ∆***ν***/*ω*_0_ [10]. The corrected foot position, ***u***_***zn***_’, is

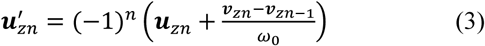

where Δ***v***_*z*_ = ***v***_*zn*_ – ***v***_*zn*-1_, and the new MoS at step *n*, ***MoS’***_*zn*_, is

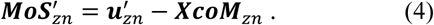

By altering the foot placement, the ***MoS***_*n*+1_, and by extension ***v***_*z*+1_, are corrected such that ***v***_*zn*-1_ = ***v***_*zn*+1_. Therefore, (3) could be written as

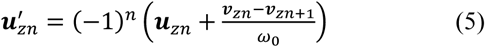

indicating a change in foot placement at step *n* can induce a change in velocity at step *n+1*. Notably, (5) is very similar to the velocity correction from (3) derived by Hof [10]. In (3) the change in foot placement should occur in the same direction as the change in velocity over the previous step. In (5), the change in foot placement should occur in the opposite direction of the intended change in velocity. From (1), (4), and (5), the change in MoS is proportional to the intended change in velocity

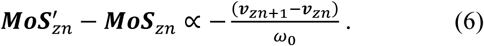

The above assumes ***MoS***_*zn*_ is the constant objective of the controller at each step and thus an arbitrary input. We can remove the constant and maintain proportionality

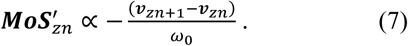

Next, we can define the centripetal acceleration of the CoM, ***a***_*c*_, as the lateral acceleration orthogonal to gravity and the direction of travel. In the lateral direction, the change in the lateral velocity of the CoM over step *n* is given by the integral

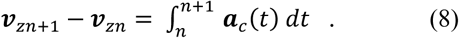

The variation in lateral MoS at heel contact can therefore be estimated using the integral of the centripetal acceleration over the following step

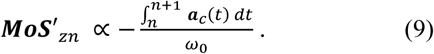

If all individuals are of relatively average stature and walking on earth, small variations in pendulum length *l* are negligible, and we can assume *ω*_0_ is a constant, and (7) can be reduced to

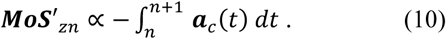

Finally, if the step time and sampling frequency are constant, as assumed for (3) by Hof [10], (9) can be reduced to the average of the acceleration over each step

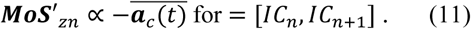

### B. Participants

Ten neurologically healthy older adults (5 Female / 5 Male) were recruited for this study. All participants provided informed written consent to participate, and all protocols were conducted in accordance with the Declaration of Helsinki and approved by the Oregon Health & Science University Institutional Review Board (IRB#15632). The study participants were an average (SD) of 72 (5.8) years of age, 169.1 (10.5) cm, and 71.5 (17.8) kg. All participants reported to be free of orthopedic and neurological impairments and medications that might affect mobility. One participant was excluded from the analysis due to a malfunctioning magnetometer throughout data collection.

### C. Experimental Protocol

All walking trials took place within a 2.5 m radius circle, marked in 45° increments around the outside (Fig 1). Within each trial, participants were instructed to pass through the center of the circle (marked in red), and then walk towards a specific colored line on the outside of the circle. For example, participants may have been given the following cue: “At your normal speed, make a slight left turn to the red line.” Thus, changing the destination color changed the turn angle. Walking trials were recorded in blocks of 10 with two trials at each turn angle (one left, one right for each of 45°, 90°, 135°, and 180°), and two straight trials per block. Three blocks were completed at a self-selected normal walking speed and three blocks performed at a self-selected fast walking speed for a total of 60 walking trials per participant.

**Fig 1.**
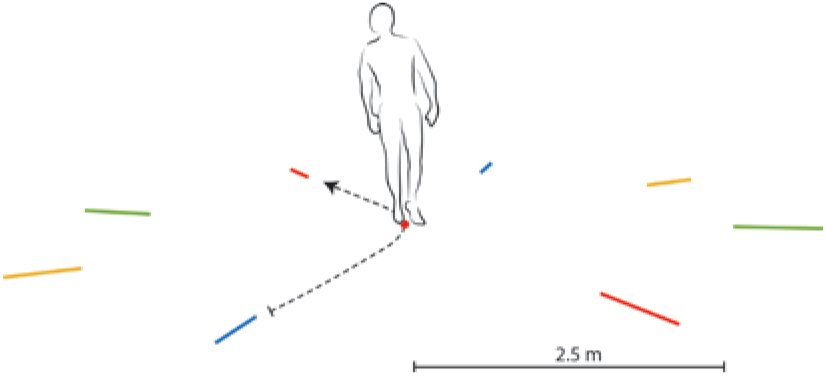
Schematic of the marked lines at 45 degree increments and center dot.

Nine inertial sensors (Opal v1, APDM Inc., Portland, OR) were placed on the following segments: forehead, sternum, lumbar spine around L3-L4, bilateral wrist, bilateral shank, and bilateral dorsum of each foot. Nine sensors were used to address other aims within the study; only three sensors, the lumbar and feet sensors, were used in this analysis. Synchronized inertial sensor data were collected at 128 Hz continuously over each block. Each block started with at least three seconds of static stance to ensure a quiet period for the sensors, but no neutral pose or specific calibration pose was collected. Additionally, all subjects were outfitted with 30 retroreflective markers in a modified Helen Hayes marker configuration (see Supplemental Figure). Markers were placed on the head (front, back, and lateral), thorax and arms (acromion, sternum, offset, lateral epicondyle of humerus, and distal radius), pelvis (sacrum, anterior superior iliac spine (ASIS)), legs (thigh, lateral epicondyle of the femur, shank, lateral and medial malleolus), and feet (1st and 5th metatarsal head, and posterior calcaneus). Optical motion capture data were collected at 120Hz (Raptor-H (8) and Osprey (4), Motion Analysis Inc., Santa Rosa, CA).

### D. Calculation of Margin of Stability

The optical motion capture data was used to calculate MoS values to validate the inertial sensor-based measures. All markers were tracked and gaps were filled using spline interpolation. All marker data were low-pass filtered using a 4^th^ order phaseless 6 Hz Butterworth filter. The instantaneous position and velocity of the whole-body CoM was estimated as the weighted average of 15 segment using kinematic data and anthropometric tables [29]. To account for the constant change in coordination frame, all data were transformed to a CoM path-of-progression reference frame aligned with gravity and the projection of the instantaneous velocity of the CoM in the transverse plane [30]. The walking speed of each trial was determined from the mean of the instantaneous CoM speed across the entire trial (including gait initiation and termination). The position of the XcoM was determined using (2), and the MoS at each point in time was determined from (1), where the lateral position of the CoP was estimated using the average of the first metatarsal and posterior calcaneus of the foot. Initial contact was defined as the maximal distance between the heel and sacrum marker in fore direction [31], and the MoS at initial contact was extracted for each step as the primary outcome. Only the MoS at initial contact was considered as previous reports have indicated the significance of this event in locomotor control [19] and fall-risk assessments [15]. All XcoM, BoS, and MoS values were oriented relative to the position of the CoM based on the vector notation in the Model Framework and the notation originally described by Hof [10]; positive MoS occured when the BoS was to the right of the XcoM, and negative MoS when the BoS was to the left of the XcoM, regardless of stance limb.

### E. Inertial Sensor Analysis

Raw inertial sensor data, including accelerometer, gyroscope, and magnetometer data were imported into MATLAB (r2018b, The Mathworks Inc., Natick, MA). Additionally, orientation estimates automatically calculated from the APDM Mobility Lab software were imported. These orientation estimates are based on Kalman filters that fuse acceleration, angular velocity, and magnetic field data to resolve quaternions between the sensor-axis and the global reference frame. For each block, the acceleration vectors at the lumbar spine were rotated to align with the global axis frame using the quaternion orientation estimates.

Subsequent analysis was completed using two different sensor alignments:

1. *Vertically Aligned Frame (VAF)*: The sensor-based local frame was rotated to align with the global vertical axis. The sensor axes were allowed to rotate about the vertical axis such that the x-axis always aligned with the direction of travel, and the z-axis aligned with the orthogonal direction. In this way, the x-z plane was always horizontal, and only yaw about the y-axis was allowed.
2. *Body-Fixed Frame (BFF)*: The sensor-based coordinate frame was fixed to the body. While initially aligned with the global frame, there was no requirement for axes to be aligned with the global frame at every instant in time throughout the trial. Sensor-based x- and z-axes may include vertical components through pitch or roll, respectively. Practically, these two alignments were obtained through either a time-varying rotation matrix (VAF) or a constant rotation matrix based on the initial alignment (BFF) between the body and global frames. Walking trials were identified and segmented into separate trials from within each block. For each walking trial, heel contacts were identified using two methods: 1) identifying peaks in the normalized frequency content above 20 Hz of the left and right foot sensors [32], and 2) using a Gaussian continuous wavelet transform of the lumbar vertical acceleration [33]. These methods use 3 sensors and 1 sensor, respectively. All steps identified using both methods were matched with steps detected from motion capture. Turns were identified within each trial using a threshold-based angular velocity algorithm (30°/s). Lumbar acceleration data were low-pass filtered using a 4^th^ order phaseless 4 Hz Butterworth filter. Centripetal acceleration at the lumbar sensor was extracted for each step (Fig 2). The acceleration was integrated between successive heel contacts based on (7). Additionally, the average acceleration between successive heel contacts was calculated based on (8). All processing steps to obtain the centripetal acceleration outcomes are shown in the Fig 3. Each of these outcomes (Integrated and Mean centripetal acceleration) were compared to the MoS at initial contact from motion capture.

**Fig 2.**
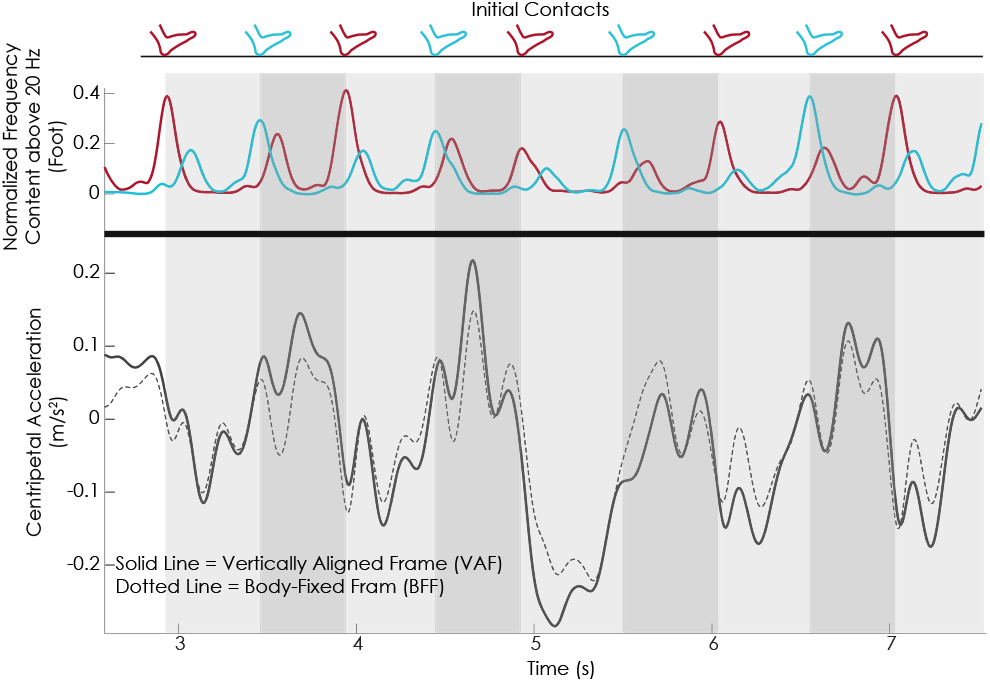
Top: Example of left (blue) and right (red) gait event detection using inertial sensors on the feet for a 90° turn. Bottom: Centripetal acceleration in vertically aligned frame (VAF, solid) and body fixed frame (BFF, dotted).

**Fig 3.**
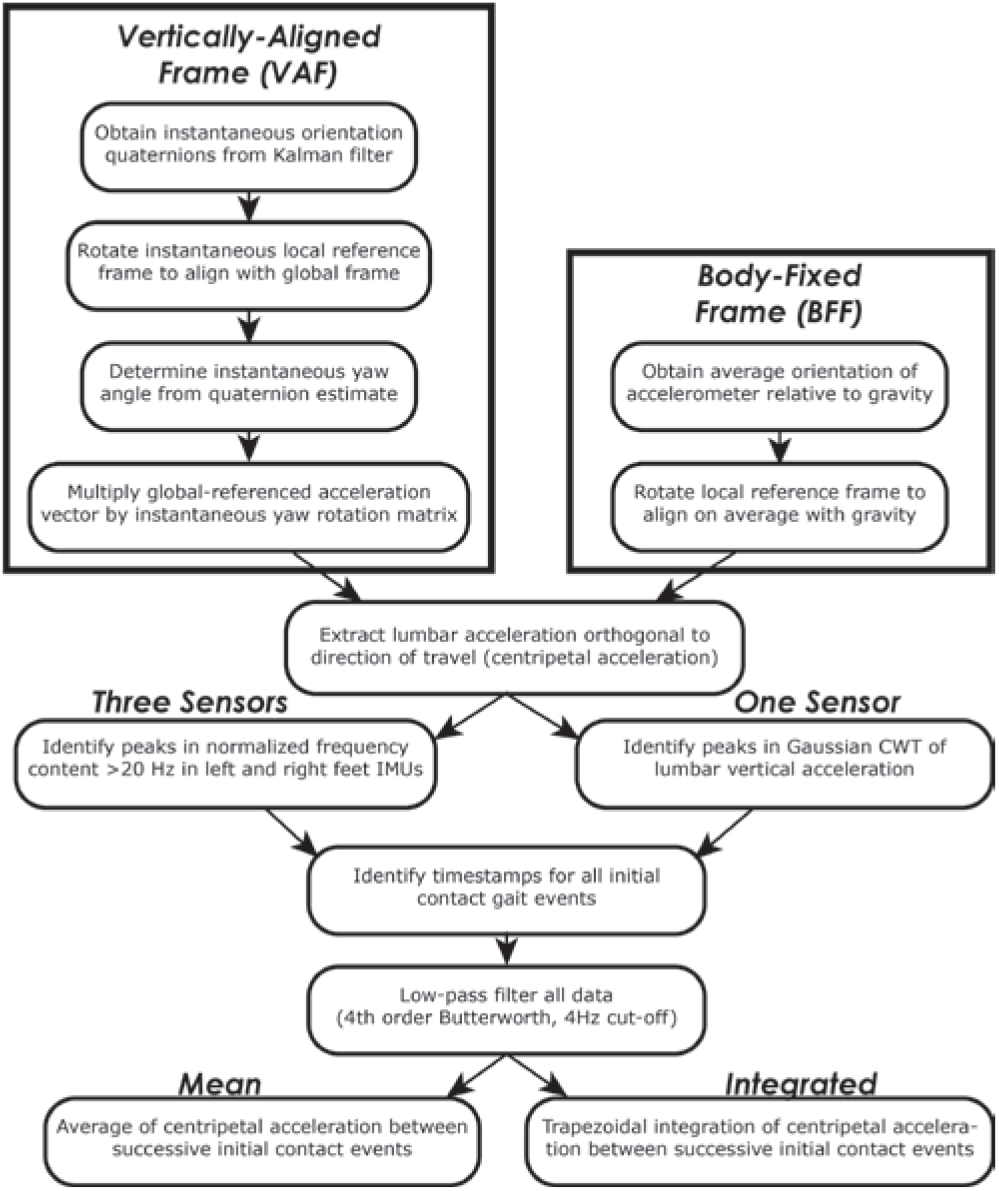
Flowchart of IMU processing pipeline to obtain VAF-, BFF-, mean, and integrated centripetal accelerations using three sensors or one sensor.

### F. Comparison Between MoS and Sensor-Based Centripetal Acceleration

To compare the level of agreement between each method of determining inertial sensor-based centripetal acceleration and the motion-capture-based MoS, linear regression models were fit using all steps from all subjects. No within-subject correction was applied, as the primary objective was to examine how well centripetal acceleration correlated with MoS regardless of subject; allowing random intercepts or slopes would not serve the intended purpose. To ensure the models were robust, a bootstrapping procedure with 10000 iterations was performed. For each model, the mean and 95% confidence interval for coefficient of determination (R^2^) was calculated from the bootstrapping. Linear regression models were also fit for each subject individually, and the range of R^2^ values was extracted to compare the between-subject performance of each method. To determine how the relationship between centripetal acceleration and MoS may change based on speed or turning angle, trials were also stratified by speed and angle, and R^2^ values were extracted from corresponding regression models.

### G. Exploring Meaningful Differences

To help guide the implementation of centripetal acceleration in future studies, we explored whether centripetal acceleration, calculated from inertial sensors, is sensitive to expected, meaningful differences. We descriptively compared the distribution of centripetal acceleration values between steps taken during straight gait and steps taken during turning. Additionally, we examined the distributions of centripetal acceleration between the inside and outside limbs during a turn, between the different turning angles, and between the different speeds. For all comparisons, centripetal acceleration values were stratified by foot to clearly illustrate the unimodal distribution per foot, and bimodal distribution when values from both feet are combined, in each condition. No statistical tests were performed; characteristics were descriptively presented to guide future metric selection and use.

## III. RESULTS

Overall, 2609 steps were included in our analyses. On average, 290 steps were included per subject (range 202-383 steps per subject) for an average of five steps per trial. The remaining steps in each trial occurred outside the volume of the motion capture cameras and therefore could not be analyzed. The average (SD) walking speeds were 1.02 (0.12) and 1.30 (0.17) m/s for normal and fast speeds, respectively.

### A. Agreement between Sensor-Based Centripetal Acceleration and Margin of Stability

Using the VAF resulted in good to excellent agreement between the sensor-based centripetal acceleration and the motion-capture based MoS (Table 1). The average centripetal acceleration over each step agreed with the MoS better than the integrated centripetal acceleration. A single lumbar-mounted sensor was near equivalent to using three sensors (lumbar, left foot, right foot) when a VAF was used in conjunction with the mean centripetal acceleration over each step – both had excellent agreement with the MoS (R^2^ = 0.73 vs. R^2^ = 0.77). The relationship between VAF-mean centripetal acceleration and MoS was consistent across subjects (Table 1, Fig 4A). The correlation between VAF-mean centripetal acceleration and lateral MoS was consistently high across turning angle using three sensors (R^2^ = 0.72, 0.82, 0.83, 0.79, and 0.71 for straight, 45°, 90°, 135°, and 180° turns, respectively), but the correlation was less consistent over turning angles using only one-sensor (R^2^ = 0.60, 0.76, 0.81, 0.75, and 0.68 for straight, 45°, 90°, 135°, and 180° turns, respectively) (Fig 4B-F). Both three- and one-sensor approaches yielded excellent relationships with lateral MoS regardless of speed; Normal and fast speeds R^2^ = 0.75 and 0.79 for three sensor VAF-mean centripetal acceleration and 0.72 and 0.74 for one sensor VAF-mean centripetal acceleration, respectively.

**Fig 4.**
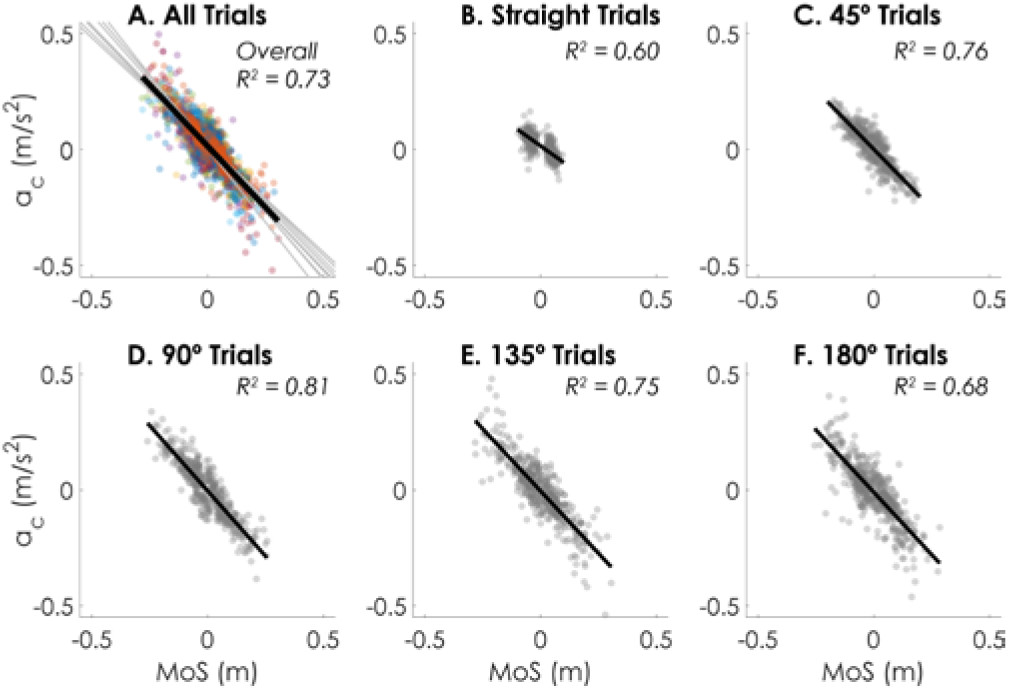
Scatter plots (A-F) between VAF-Mean centripetal acceleration using a single sensor and lateral MoS. A. All steps from all trials are included, with different colored scatters indicating different subjects. Thin gray lines indicate linear fits within each subject. The thick black line indicates the linear fit across all subjects and trials. B-F. Scatter plots for trials of specific turning angles, with all subjects plotted together and overall linear fits depicted with thick black lines.

**TABLE I.** Correlation between Mos and IMU-Based Centripetal Acceleration

| | $R^2$ | 95% CI | RMSE | $R^2$ (Subject Range) |
| --- | --- | --- | --- | --- |
| **Three Sensors** |  |  |  |  |
| VAF - Integrated | 0.75 | 0.73-0.76 | 0.04 | 0.61-0.87 |
| VAF - Mean | 0.77 | 0.75-0.79 | 0.04 | 0.64-0.87 |
| BFF - Integrated | 0.44 | 0.40-0.47 | 0.06 | 0.10-0.60 |
| BFF - Mean | 0.43 | 0.39-0.47 | 0.06 | 0.11-0.72 |
| **One Sensor** |  |  |  |  |
| VAF - Integrated | 0.66 | 0.63-0.69 | 0.05 | 0.41-0.82 |
| VAF - Mean | 0.73 | 0.71-0.75 | 0.04 | 0.59-0.84 |
| BFF - Integrated | 0.35 | 0.31-0.38 | 0.07 | 0.05-0.68 |
| BFF - Mean | 0.37 | 0.33-0.41 | 0.07 | 0.09-0.74 |

VAF – Vertically-aligned frame; BFF – Body-fixed frame

### B. Straight Gait versus Turning

Comparing straight walking and turning, straight walking had a much tighter distribution of centripetal acceleration, centered at 0, compared to turning (Fig 5), agreeing with the expectation that centripetal acceleration is minimal during straight travel.

**Fig 5.**
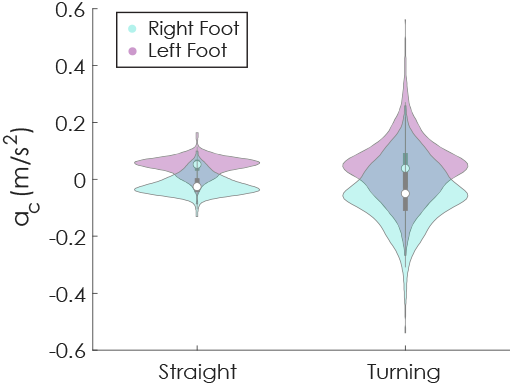
Violin plot depicting the distributions of centripetal acceleration values during straight and turning trials, stratified by stance limb. Note that the stratification is by trial, and therefore turning trials include all steps in the trial, including straight steps.

### C. Inside versus Outside Limb

During left and right turns, the distributions of the average centripetal acceleration when the inside foot was in stance (left foot during left turns, right foot during right turns) were close to zero, and inside limb left and right foot plots overlapped more than during straight walking. Distributions of the outside limb were centered away from zero and skewed (Fig 6).

**Fig 6.**
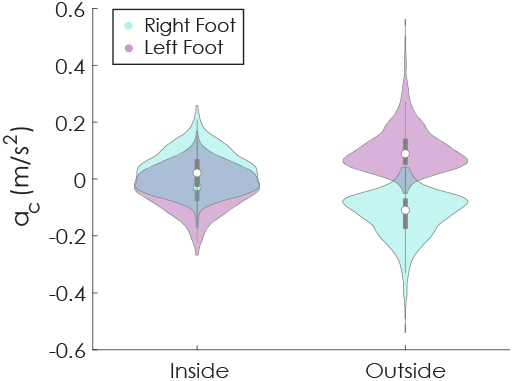
Violin plot depicting the distributions of centripetal acceleration between the inside and outside stance limb, stratified by foot.

### D. Difference between Turning Angles

While sharper turning angles tended to widen distributions and increase the variance compared to shallower turning angles, this trend was only truly noted when comparing 45 degree turns to sharper turns (Fig 7). Note that all turns had a concentration of accelerations similar to straight gait due to the protocol.

**Fig 7.**
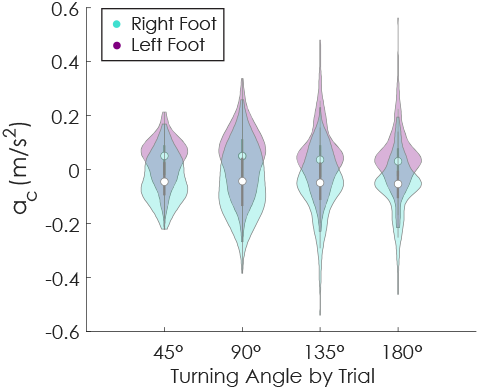
Violin plot depicting the distribution of centripetal acceleration at each turning angle. Qualitative differences can be noted as the turning angle increases, with more extreme values in centripetal accelerations. Note that all turning angles have a concentration resembling straight gait due to each trial including steps towards and away from the center dot.

### E. Difference between Speeds

Speed primarily affected the centripetal acceleration on the inside limb of the turn. During fast trials, greater centripetal acceleration magnitudes were evident on the inside limb compared to normal walking trials (Fig 8).

**Fig 8.**
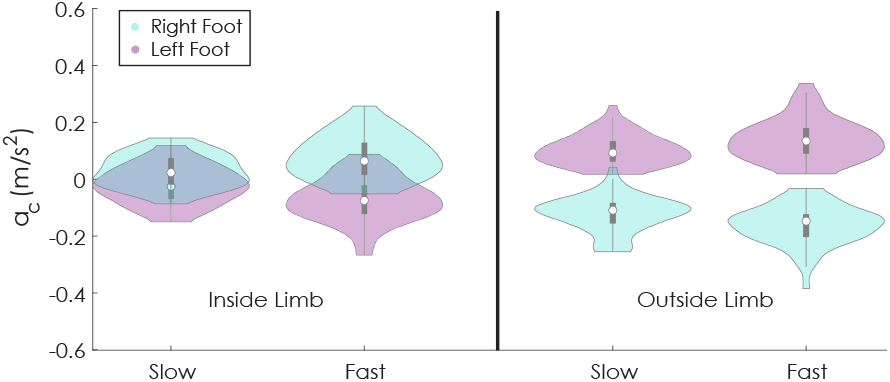
Violin plots for 90 degree turns only, stratified by left and right foot and inside and outside limb. Speed primarily changed the centripetal acceleration of the inside limb.

## IV. DISCUSSION

Centripetal acceleration calculated from inertial sensors on the feet and the lumbar spine was able to estimate lateral MoS during walking and turning. Notably, the relationship between average centripetal acceleration and MoS was consistent and strong across all subjects without the need for a subject-specific correction for anthropometry (Fig 4). However, the validity of this estimation required using a VAF and the average of the centripetal acceleration over the following step. Restricting the analysis to a single sensor on the lumbar spine resulted in negligible decrements in performance. The largest difference between three-sensor and one-sensor approaches appeared during straight walking trials (R^2^ = 0.72 vs. 0.60, respectively), suggesting a three-sensor approach may be necessary when investigating straight gait. Nevertheless, these results suggest that MoS during combined walking and turning may be estimated using only a single inertial sensor located around the waist with the caveat that a method for determining which limb is in stance is recommended for the single sensor solution.

Interestingly, the best agreement between the centripetal acceleration and lateral MoS was found by averaging, rather than integrating, the centripetal acceleration over each step. This result was curious because the construct of averaging relied on assumptions of constant step time. The improved performance of averaging, compared to integrating, can most likely be attributed to a lack of precision in our gait event detection and error propagation from integration. As seen in Fig 2, step transitions correspond to large shifts in the centripetal acceleration. Small temporal errors in step detection, therefore, are more likely to compound when integrated than when averaged. The decrease in performance of the single-sensor, integrated acceleration algorithm provides further support along this line as the precision of gait event detection decreases when using a single lumbar-mounted sensor compared to sensors on the lower extremities [34]. Additionally, random error propagates when integrated (often called drift), while averaging reduces error. Integrating, while based on less model assumptions, is therefore more susceptible to random error in accelerometer measurements. While step time is not constant, variation in step time is small in healthy adults (<3-5%) [35], and an approximation of constant step time is not unreasonable in this population. Combined, these factors suggest averaging the acceleration over each step, even if assumptions are not strictly met, is more robust against random gait event detection and measurement error than an integration-based approach.

### A. Limitations and Guidelines for Implementation

While the average centripetal acceleration over each step was consistent and valid across subjects and sensitive to different conditions, underlying assumptions and limitations must be considered when using centripetal acceleration from inertial sensors as a correlate for lateral MoS. Specifically:

1. *Correctly identifying left and right foot contacts may be problematic using only a single sensor*. Previous methods have relied on using the lateral acceleration or angular velocities to determine the stance limb using a lumbar-mounted inertial sensor [33]. However, angular-velocity-based methods are not viable during non-straight gait, where roll and yaw angular velocities are strongly influenced by the turn. In these cases, left-right stance limbs are assumed based on alternating steps within a pair. While this is generally a robust assumption, it is not always true, particularly for sharp turns and in individuals with severe gait impairments. For this reason, if a primary outcome is dependent on identifying MoS on each foot, we recommend using the three-sensor approach until a validated method emerges addressing this problem. While results may be meaningful without stratifying by stance limb, interpretations should carefully consider the underlying bimodal distributions.
2. *Step-to-step based centripetal acceleration correlates of lateral MoS may not be robust for comparisons with small effects*. Inertial sensors matched motion-capture-based MoS with R^2^ values exceeding 0.7 and, on average, were very consistent. However, ~ 25% of variance remained unexplained. As noted in Fig 4, many points fall along the correlation line of best fit, but some do not. Therefore, it is advisable to use aggregate summary statistics, rather than individual maximum or minimum values, to compare conditions. Further work should validate the accuracy and reliability of centripetal acceleration for individual perturbation recovery steps.
3. *Centripetal acceleration may not be reliable in scenarios with external forces (e.g., perturbations)*. Based on our model framework, the average centripetal acceleration over one step is dependent on the desired change in CoM velocity at initial contact. Therefore, there is a time lag that must be considered when external forces are applied. For instance, a lateral impulse *J* applied to the CoM during stance will change the centripetal acceleration of the CoM, but will not retroactively adjust the MoS at the initial contact preceding that stance. In this case, the average centripetal acceleration over stance will differ from the MoS by *J*/*m*, where *m* is the mass of the individual.
4. *Mean centripetal acceleration may not be useful in some comparisons of continuous monitoring*. As noted in Fig 5, the distribution of centripetal acceleration is distinct between walking and turning. However, comparing only mean values does not capture the full picture; the spread of the distribution is the most apparent difference between walking and turning. As daily walking is a continuous mixture of straight and turning steps, examining the variability of centripetal acceleration may be advisable considering the underlying bimodal distributions.
5. *Reliance on the VAF requires robust sensor fusion algorithms and stable magnetometer estimates*. Average centripetal acceleration only related to MoS when centripetal acceleration was confined within the global horizontal plane (VAF). To achieve this VAF, continuous estimates of the lumbar sensor orientation had to be resolved by fusing accelerometer, gyroscope, and magnetometer data. In unknown environments, changes in the local magnetic field may influence the magnetometer reading and alter the alignment of the VAF. Uses of centripetal acceleration as a correlate for MoS should consider using sensor fusion algorithms that are robust to environmentally-induced magnetometer changes.
6. *Validity in pathological populations has not been established*. Only healthy older adults were tested here. While the long-term utility of this approach may include continuous monitoring of pathological populations, it is unclear whether the centripetal acceleration will maintain its consistent relationship. Populations with short, shuffling steps may pose particular problems associated with gait event detection.
7. *Ignoring the eigenfrequency may have more significant effects in different populations*. Our sample of adults was relatively homogenous in stature. It is possible that the effects of eigenfrequency, which were ignored in this analysis due to the small variance, may need to be accounted for in populations with widely varying stature (e.g., children vs. adults).

## V. CONCLUSIONS

Inertial sensors can provide reliable and consistent measures of the centripetal acceleration of the CoM that correlate with the lateral MoS. While the best results were obtained using an inertial sensor on each foot and one on the lumbar region of the spine, output from a single sensor on the waist is also capable of providing valid and robust estimates of the lateral MoS given knowledge of stance foot at time of calculation. It is possible to obtain reliable MoS estimates using only a few inertial sensors, but future validation may be required during free-living walking in community settings using this approach. Limitations and assumptions prompt future work.

## ACKNOWLEDGMENT

The authors thank Spencer Smith, Graham Harker, and Georgeann Booth, Maddy Dunn, and Grace McBarron for assisting with data collection and subject recruitment.

