## Supplemental Figure for "Inertial sensor-based centripetal acceleration as a correlate for lateral margin of stability during walking and turning"

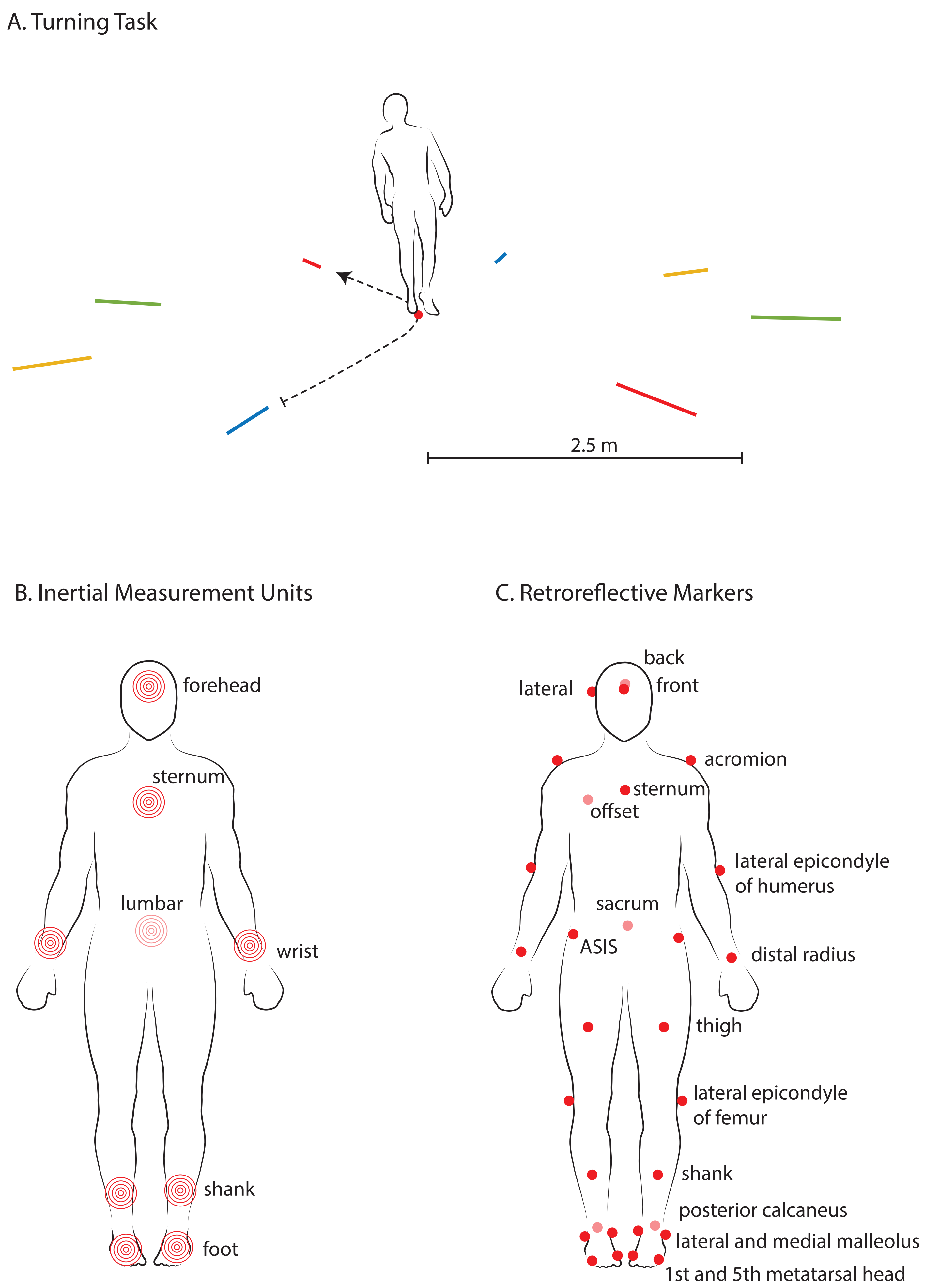


Supplemental Figure 1. Experimental overview A.) Depiction of the protocol with marked lines at 45 degree increments and center dot. B.) Illustrative depiction of the placement of each inertial measurement unit on the body. C.) Illustrative depiction of the placement of retroreflective markers on the body for optical motion capture.
